# Infrared Spectroscopy for Structure Analysis of Protein Inclusion Bodies

**DOI:** 10.1101/2022.07.20.500777

**Authors:** Andreas Schwaighofer, Bernhard Lendl

**Author notes:** Correspondence and requests for materials should be addressed to: Andreas Schwaighofer, TU Wien, Research Group Process Analysis, Institute of Chemical Technologies and Analytics, Getreidemarkt 9/164, A 1060 Vienna, Austria.

## Abstract

Infrared (IR) spectroscopy is a widely used technique for evaluation of protein secondary structure. In this chapter, we focus on the application of this analytical technique for analysis of inclusion bodies. After a general introduction to protein analysis by IR spectroscopy different approaches for spectra acquisition, data processing and secondary structure evaluation are presented.

## 1. Introduction

Mid-Infrared (IR) spectroscopy is a powerful tool to study the structure of proteins, and as such also inclusion bodies (IBs) [1, 2]. Compared to other spectroscopic methods for protein analysis such as circular dichroism (CD) spectroscopy, IR spectroscopy is particularly sensitive to aggregated protein structures as they commonly occur in IBs. Furthermore, when employing IR spectroscopy less care needs to be taken in regard to turbidity and salt concentration of the sample [3].

IR spectroscopy detects the vibrational transitions of functional groups in the spectral region of 400-4000 cm^-1^ (25-2.5 µm). The most prominent absorption bands of proteins are the amide I band (1600-1700 cm^-1^ - originating from the C=O stretching and N-H in-phase bending vibration of the amide group) and the amide II band (1500-1600 cm^-1^ - arising from N-H bending and C-N stretching vibrations). The amide I band has been recognized to be most sensitive to the secondary structure. Its sensitivity to individual secondary structure elements originates in differing patterns of hydrogen bonding, dipole-dipole interactions, and geometric orientations in the α-helices, β-sheets, turns, and disordered structures. These varying interactions induce different frequencies of the C=O vibrations and consequently in the characteristic band maxima and shapes. Accordingly, the individual secondary structure elements absorb in specific sections of the amide I band [4], as summarized in Table 1.

**Table 1.** Assignment of amide I band positions to secondary structure [4].

| Secondary structure | Band positions in H<sub>2</sub>O / $\text{cm}^{-1}$ | |
| --- | --- | --- |
|  | Average | Range |
| α-helix | 1654 | 1648-1657 |
| β-sheet | 1633 | 1623-1641 |
|  | 1684 | 1674-1695 |
| Turns | 1672 | 1662-1686 |
| Disordered | 1654 | 1642-1657 |

Absorption bands of α-helices highly overlap with disordered structures, thus it is generally difficult to distinguish these structure elements by IR spectroscopy. In contrast, the main absorption band of native β-sheet secondary structure appears isolated at the low-wavenumber flank of the amide I band and is rather undisturbed from absorptions of other secondary structures.

Protein aggregates in inclusion bodies feature a high content of intermolecular β-sheet structures. For those, the absorption band is uniquely shifted to even lower wavenumber regions (1615-1625 cm^-1^) due to its highly repeated structure elements. β-sheet structures also often show a distinctive second, weaker band at the high-wavenumber end. Furthermore, β-sheet structures exhibit particularly high molar absorptivity among the individual secondary structure elements [5]. Due to the isolated position of the related absorption band and the high sensitivity to β-sheet structures, IR spectroscopy is most suited for analysis of protein aggregates and IBs. Thus, IR spectroscopy was successfully employed to address different questions of IB analysis, including structure analysis [6-11], IB quantification [12], monitoring of IB formation kinetics [13], protein refolding [14] and IB activity [15].

In the following sections a guideline is given on how to record high quality IR spectra of IB samples combined with a workflow on spectra processing and secondary structure analysis.

## 2. Materials

### 2.1. Basic Lab Equipment and Chemicals

1. Lab Centrifuge (e.g., MiniSpin by Eppendorf)
2. Centrifugal filters for buffer exchange
3. Ultrapure water, optional: deuterated water (D_2_O)
4. Disodium hydrogen phosphate (Na_2_HPO_4_), monosodium dihydrogen phosphate (NaH_2_PO_4_), or PBS (phosphate buffered saline) tablets
5. 1% sodium dodecyl sulfate (SDS) solution and 70% ethanol (C_2_H_5_OH) for cleaning of transmission cell or ATR accessory

### 2.2. IR Instrumentation and Accessories

#### 2.2.1. IR spectrometer

1. FTIR Spectrometer There are multiple manufacturers for (Fourier-transform IR) FTIR spectrometers, including Bruker Optics, Thermo Fisher Scientific, Perkin Elmer. Each producer provides proprietary software for instrument control and spectra acquisition.
2. Laser-based IR spectrometer Commercial laser-based IR spectrometers are available from Daylight Solutions (ChemDetect, Culpeo) and RedshiftBio (AQS^3^pro, Apollo). Due to the higher spectral power densities of laser light sources, with laser-based IR instruments higher path lengths can be used for liquid-phase transmission measurements and consequently enable significantly more robust sample handling.

#### 2.2.2. Transmission cell

For liquid-phase transmission measurements, demountable flow cells are used consisting of a mount, cell windows and a spacer. Most commonly used window materials are zinc selenide (ZnSe) and calcium fluoride (CaF_2_). Both are hard materials and resistant to most solvents, however should only be used in moderate pH conditions (pH between 5 and 9). Furthermore, complexing agents, such as ammonia and EDTA, may erode the surface of ZnSe due to the formation of complexes with zinc. ZnSe has a wider transmission range in the low wavenumber region (cutoff 500 cm^-1^ vs cutoff 1000 cm^-1^ of CaF_2_), which however for protein analysis is neglible. An important advantage of CaF_2_ over ZnSe is its lower refractive index of 1.4 (vs. 2.4 for ZnSe) which matches better to aqueous solutions. As a consequence, interference fringes leading to excess noise in the recorded spectra are less of a problem when using CaF_2_ windows.

For analysis of the protein amide I band, spacers with a thickness of ∼10 µm are employed for FTIR spectroscopy and ∼25-35 µm for laser-based IR spectroscopy.

Demountable transmission cell mounts are available by e.g., Perkin Elmer or Specac. Cell windows can be purchased at e.g., Crystran or Korth Kristalle.

#### 2.2.3. ATR accessory

For (attenuated total reflection) ATR measurements, accessories for FTIR spectrometers are commercially available. Versatile single-bounce ATR accessories with a diamond reflecting element are available from multiple manufacturers (e.g., Platinum ATR by Bruker Optics, Golden Gate ATR by Specac).

## 3. Methods

### 3.1. IR spectra acquisition

To obtain an IR absorbance spectrum, two intensity (in FTIR spectroscopy also often called single channel) spectra need to be recorded. The reference spectrum, 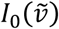, is recorded of the sample matrix. This can be the solvent (buffer, pure water) in case of liquid-phase samples or air/plain ATR crystal for analysis of dried samples by ATR spectroscopy. The sample spectrum, 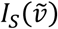, is recorded of the analyte in its matrix. From these intensity spectra, the absorbance spectrum is calculated by

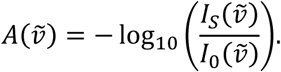

Consequently, all differences between the reference (matrix) and sample (matrix+analyte) measurements are represented in the calculated absorbance spectra. This relates not only to the presence of the analyte in the sample, but also to any changes that occur in the matrix between the reference and sample measurement. For this reason, great care must be taken that the matrix composition (e.g. salt concentrations, buffer concentration, pH value) as well as the measurement conditions (e.g. temperature) do not change between the two measurements. This also applies to the humidity within the IR instrument, however, water vapor absorption lines can be straight-forwardly corrected as outlined in section 3.2.1.

A major advantage of IR spectroscopy is the possibility to record spectra from different sample states. IB samples can be investigated as aqueous solution, hydrated films or even dry solids. The two typically used sampling techniques are transmission and ATR, see Figure 1. For transmission measurements, the liquid sample is inserted into a transmission cell by a syringe or pump. Depending on the type of tubings and the dead volume of the cell, 100-500 µL of sample solution are needed. In contrast to transmission spectroscopy, where the sample is inserted in a measuring cell, for ATR measurements the sample is placed on the surface of an optically dense ATR element. In this measurement technique, the infrared light is reflected at the contact surface of the ATR element and the sample, whereby a small part of the IR light penetrates the sample as an evanescent wave and can interact with the sample. Here, a drop of sample is sufficient to obtain an IR spectrum.

**Fig. 1.**
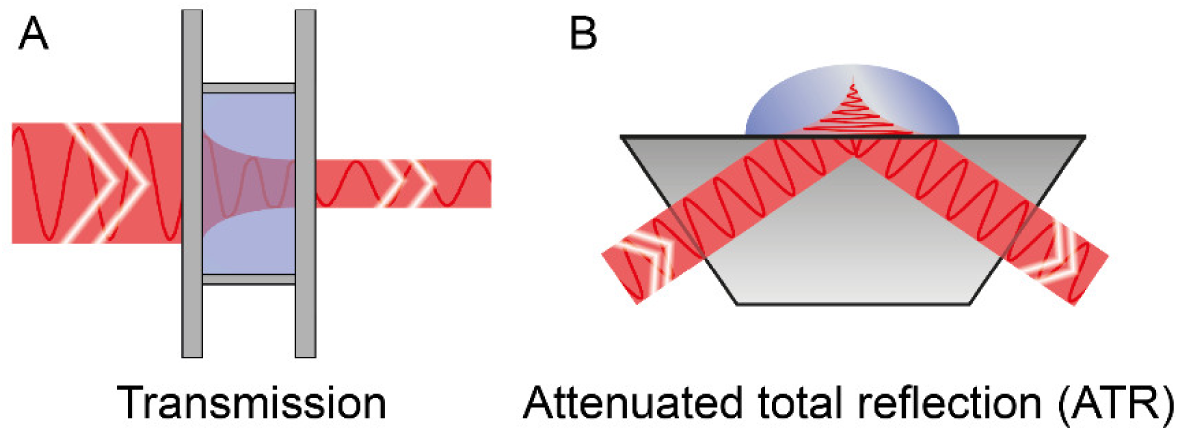
IR sampling methods. **(A)** Transmission. **(B)** Attenuated total reflection (ATR).

A pronounced advantage of ATR spectroscopy is the low required time and effort in terms of sample preparation and method optimization, as the sample merely needs to be placed on the ATR crystal. In contrast, sample handling with transmission measurements may be rather laborious as thin Teflon spacers are extremely susceptible to electrostatic charging thus making the assembly of tight flow cells with ∼10 µm path length difficult.

Transmission spectroscopy offers advantages in terms of limit of detection, in case only low analyte concentrations are available or quantitative analysis is desired. In absorption spectroscopic techniques, quantitation is performed according to the Beer-Lambert”s law

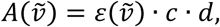

where A denotes the measured absorbance, 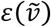 refers to the molar decadic absorption coefficient, c is the analyte concentration and d signifies the path length. Consequently, higher absorbance signals can be obtained by employing a larger path length. For transmission measurements in aqueous solution, the path length is approx. 10 µm. In ATR spectroscopy, the path length depends on the wavelength, the angle of incidence and the refractive index of the ATR element, however for commercial single-reflection accessories, it typically amounts to 0.5-2 µm [16]. However, multibounce ATR accessories can be employed to increase the measured signal and by that the sensitivity.

An alternative approach to increase the sensitivity for transmission measurements is to perform IR analysis in deuterated solution. D_2_O mostly has similar physical and chemical properties when compared to regular water. For IR spectroscopy, this solvent offers the advantage that the DOD-bending band is positioned at approx. 1200 cm^-1^, providing a region of relatively low solvent absorption between 1500 and 1800 cm^-1^. The shift of the intense solvent absorption bands to lower wavenumbers of the solvent enables using longer path lengths for IR measurements in the amide region. Typical transmission path lengths for secondary structure analysis by FTIR spectroscopy is in the range of 100 µm. This increase of the path length compared to aqueous measurements is accompanied by an increase of the sensitivity as outlined above. The shape of protein spectra recorded in deuterated solution appears different because the amide II band (predominantly originating from N-H vibrations) is shifted from 1550 to 1450 cm^-1^. The amide I band is mainly composed of C=O vibrations, thus there occurs only a small shift to lower wavenumbers by 5-10 cm^-1^. Consequently, also the spectral regions associated to individual secondary structure elements are shifted to lower wavenumbers as compared to Table 1 [4].

#### 3.1.1. Common FTIR spectra acquisition parameters

1. Recorded spectral range: 4000-800 cm^-1^.
2. Spectral resolution: 2 cm^-1^.
3. Accumulated scans: 128.
4. Interferogram acquisition mode: Double-sided, forward-backward.
5. Apodization function: Blackman-Harris 3-term.
6. Zero filling factor: 2.

#### 3.1.2. Sample preparation

Some buffer substances (e.g. Tris, citrate, acetate) as well as stabilizing agents (e.g. glycerol) or chaotropic substances (e.g. guanidine hydrochloride, urea) show absorption bands overlapping with the protein amide I and amide II bands. Consequently, it is beneficial to dialyze the protein solution against pure water or phosphate buffer to remove unwanted components.

When performing liquid-phase measurements, the IB concentrations should be >3 mg/mL for transmission measurements and >5 mg/mL for ATR measurements. In case the concentration is significantly lower, the IB sample should be concentrated, after or concurrent with purification steps. For IR measurements of IBs dried on an ATR crystal, the concentration can be lower.

#### 3.1.3. (Micro)Spectroscopy of dried IBs in transmission

1. Centrifuge IBs and remove supernatant.
2. Resuspend IB pellet in ∼30 µL of distilled water.
3. Take a reference spectrum of blank ZnSe/CaF_2_ window.
4. Deposit ∼15 µL of the sample suspension on the ZnSe or CaF_2_ window and let it dry for 30 min. The deposited volume defines the spot size area.
5. Take a sample spectrum.
6. Clean window with an excess of water and if necessary, with 1% SDS or 70% ethanol. For thorough cleaning, the windows can also be treated in an ultrasonic bath.

#### 3.1.4. ATR-IR spectroscopy of dried IBs

1. Centrifuge IBs and remove supernatant.
2. Resuspend IB pellet in ∼30 µL of distilled water.
3. Take a reference spectrum of blank ATR crystal.
4. Deposit ∼20 µL of the sample suspension on the ATR crystal and let it dry for 30 min. The surface area of the ATR crystal should be completely covered.
5. Take a sample spectrum of dried IB film.
6. Clean ATR crystal with an excess of water and if necessary, with 1% SDS or 70% ethanol.

#### 3.1.5. Transmission spectroscopy of IBs in aqueous solution

Because of the high background absorption of water in the protein amide I region, short transmission path lengths in the range of 10 µm are required to enable acquisition of high-quality IR spectra. Aggregated IBs can span a wide size variety from small oligomers to micron-sized aggregates and may thus impair sample handling und lead to clogging of narrow transmission cells. If this is the case for a particular sample, then a larger transmission path length can be used employing a laser-based spectrometer [8, 9, 17, 18], if available, or ATR measurements are preferrable.

1. Prepare a transmission cell with path length ∼10 µm for FTIR spectroscopy or ∼25 µm for laser-based spectroscopy.
2. Record reference spectrum of flow cell filled with sample matrix i.e water or buffer.
3. Prepare IB sample as outlined in Section 3.1.2, if necessary.
4. Introduce aqueous sample into transmission cell by syringe or pump.
5. Record sample spectrum of flow cell filled with sample solution.
6. Clean the transmission cell with an excess of water/buffer and if necessary, with 1% SDS or 70% ethanol.

#### 3.1.6. ATR-IR spectroscopy of IBs in aqueous solution

In case the aggregated structures of the IB sample are too large to be ruggedly measured with transmission cells, FTIR analysis in solution can also be performed by the ATR technique.

1. Deposit ∼20 µL of the sample matrix/solvent on the ATR crystal. The surface area of the ATR crystal should be completely covered.
2. Take a reference spectrum of the sample matrix/solvent.
3. Prepare IB sample as outlined in Section 3.1.2, if necessary, and deposit ∼20 µL on the ATR crystal.
4. Take a sample spectrum of the IB sample.
5. Clean ATR crystal with an excess of water and if necessary, with 1% SDS or 70% ethanol.

#### 3.1.7. Transmission spectroscopy of IBs in D_*2*_*O solution*

Transmission measurements of IB samples can be performed also in deuterated buffer solution, as outlined above. Due to the higher transmission path length up to 100 µm, lower IB concentration of >0.5 mg/mL can be measured. Sample transfer to deuterated solution can be achieved by two approaches:

1. Dialysis of the IBs against pure D_2_O or phosphate buffer prepared in D_2_O.
2. Lyophilization of aqueous IB sample to dryness. Then reconstitution of sample in small amount of D_2_O and again lyophilization to dryness. Finally, reconstitute IB sample in pure D_2_O or phosphate buffer prepared in D_2_O.

The thereby obtained samples can be analyzed as follows:

1. Assemble transmission cell with path length ∼100 µm for FTIR spectroscopy and ∼400 µm for laser-based spectroscopy.
2. Record reference spectrum of flow cell filled with sample matrix i.e. D_2_O or deuterated buffer.
3. Introduce liquid-phase sample into transmission cell by syringe or pump.
4. Record sample spectrum of flow cell filled with sample solution.
5. Clean the transmission cell with an excess of D_2_O/deuterated buffer and if necessary, with ethanol.

### 3.2. IR spectra processing

Spectra processing is an important step to prepare experimental results for ensuing spectra analysis and interpretation. The presented processing steps can be conveniently performed by (often proprietary) software provided by spectrometer manufacturers, but also by general data analysis software such as OriginLab Origin, MATLAB or python.

#### 3.2.1. Correction for atmospheric water vapor

In the spectral region of the protein amide I + II bands, there also occur absorption bands of the rotational transitions of water vapor, which is ubiquitously present in our ambient environment. In case the humidity changes between acquisition of the reference and sample intensity spectrum, water vapor bands appear in the IR spectrum in form of sharp absorption lines. For this reason, the time period between these two measurements should be kept short. However, for long-term experiments variations in ambient humidity often cannot be avoided. Experimental measures to improve the stability include purging the spectrometer with dry air or nitrogen. If this is not possible, drying agent can be placed in the instrument.

In case water vapor absorption bands cannot be avoided experimentally, they can be corrected from the recorded spectra by several methods. For these computational approaches, it is advantageous to record the spectra at relatively high resolution (2 cm^-1^) to benefit from the width difference between the water vapor and liquid sample bands. The most straight forward approach for correction is direct subtraction of a reference water vapor spectrum multiplied with a scaling factor. This coefficient can be either determined visually or computationally. For manual correction, the water vapor spectrum is subtracted from the sample spectrum until the spectral region between 1700 and 1800 cm^-1^ appears featureless. The corrected spectra can be finally smoothed to a resolution of 4 cm^-1^, which is still high enough for protein structure analysis. Fig. 2 depicts IR spectra of water vapor as well as an IB sample before and after correction. Procedures describing more complex approaches for atmospheric water vapor correction can be found elsewhere [19].

**Fig. 2.**
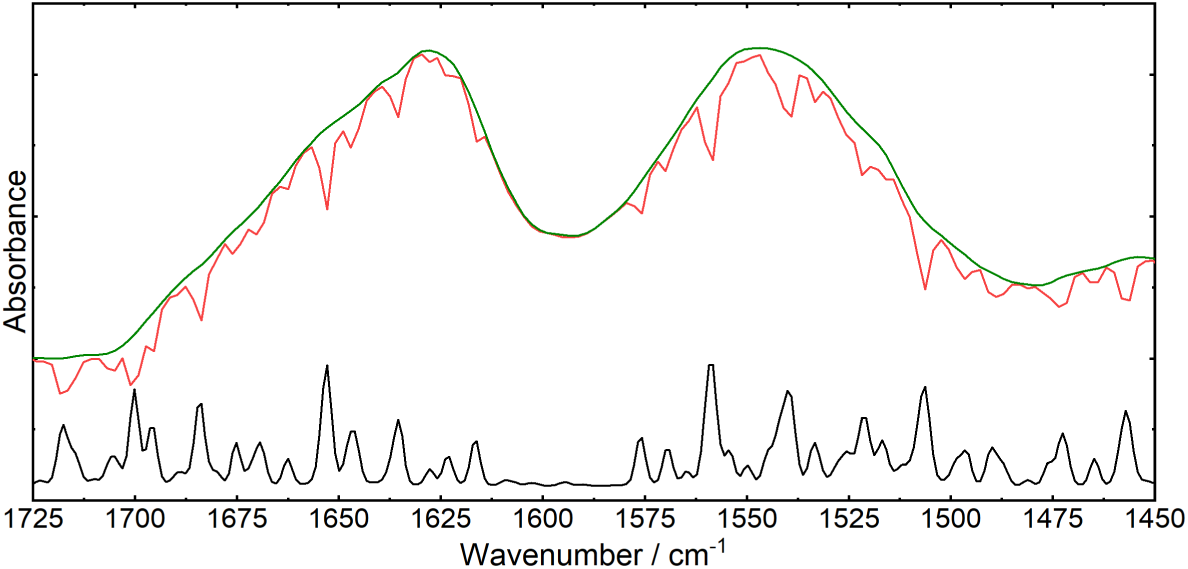
Absorbance spectrum of an IB sample recorded in the presence of water vapor (red). Corrected absorbance spectrum (green). Absorbance spectrum of atmospheric water vapor recorded at a spectral resolution of 2 cm^-1^ (black).

#### 3.2.2. Difference spectra

Difference spectra are calculated with the aim to simplify IR spectrum interpretation by guiding the focus on the spectral changes between two or multiple IR spectra. By calculation of difference spectra 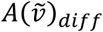

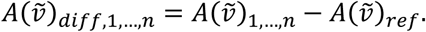

with 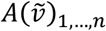 as the spectra of interest and 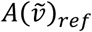 as the reference spectrum, the spectral differences to the reference spectrum are highlighted by eliminating bands with the same shape, wavenumber positions and intensities. Since this kind of spectral differences can be also unintentionally introduced by varying measurement conditions such as e.g. temperature, great care must be taken during experimental data acquisition. This processing step is often applied when monitoring dynamic processes. The initial spectrum is taken as reference und the resulting difference spectra contain only the bands that emerge during the investigated process [20]. Furthermore, difference spectroscopy is also a useful tool to highlight the spectral differences between static spectra. Fig. 3A shows the absorbance spectra of a correctly folded protein and an IB sample of the same protein. Fig. 3B shows the difference spectra calculated with the protein sample as a reference. Here, the spectral differences are emphasized and the higher content of aggregated protein structure (1625 cm^-1^) in the IB sample is highlighted.

**Fig. 3.**
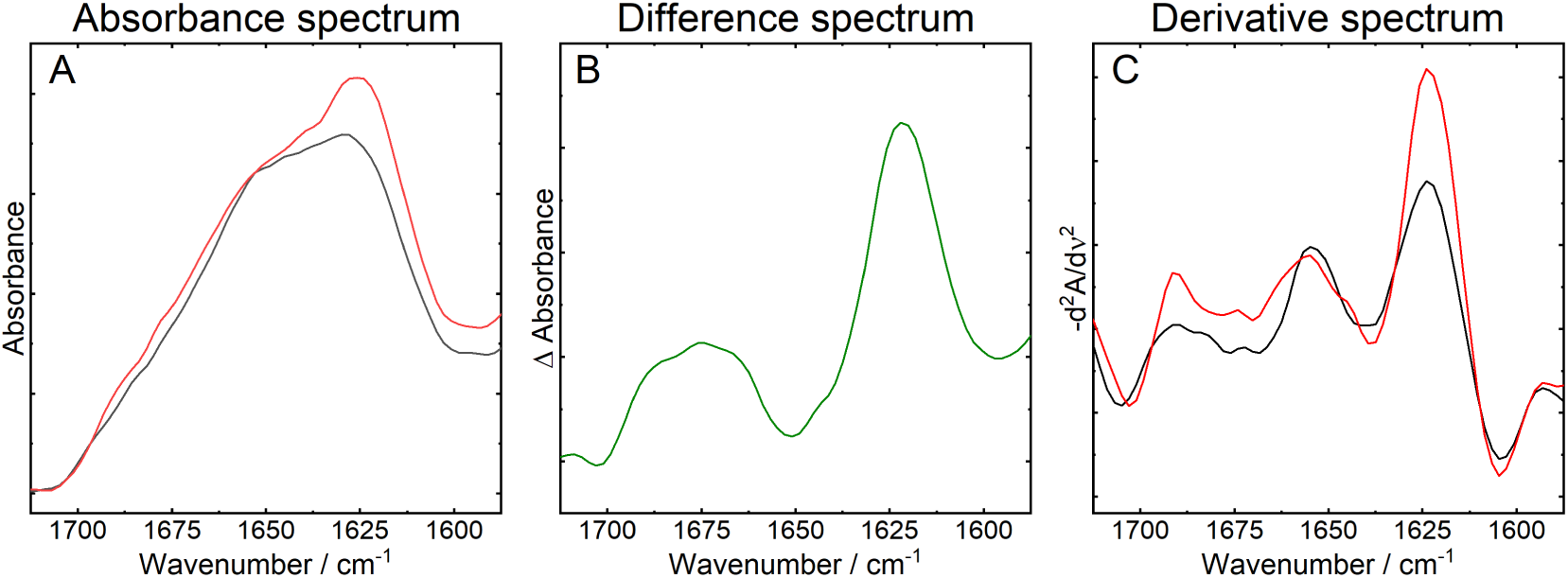
**(A)** Absorbance spectrum of a correctly folded protein sample (black) and an IB sample (red). **(B)** Difference spectrum of the IB sample with the correctly folded protein sample as a reference. **(C)** Inverted second derivative spectrum of the correctly folded protein sample (black) and an IB sample (red).

#### 3.2.3. Second derivative spectra

Analysis of derivative spectra is a frequently applied approach for resolution enhancement. By calculation of derivatives of the experimentally recorded absorption spectra, the widths of the IR bands are narrowed which in turn leads to a seemingly improvement of the spectral resolution and enables deconvolution of overlapping bands. For straight forward analysis, spectroscopists most often employ the inverse second derivative spectrum for analysis, because even-ordered derivatives have their minima/maxima at the same wavenumber positions as the original absorbance spectrum. As in the case of the second derivative, there occurs a minimum and thus the derivative spectrum is then inversed to obtain positive bands. An important feature of derivative spectra is that the quantitative properties of the original spectrum are preserved, thus they can be employed for quantitation. Calculation of derivative spectra significantly decreases the signal-to-noise-ratio, therefore this calculation should be accompanied by a smoothing step. Fig. 3C shows the inverted second derivative spectra of the absorbance spectra depicted in Fig. 3A. In these spectra, the band maxima at 1625 and 1655 cm^-1^ as well as the shoulder at 1690 cm^-1^ become particularly prominent.

### 3.3. Secondary structure analysis by curve fitting

Curve fitting of the amide I band is the most commonly used method to estimate the relative portions of the individual secondary structure elements of a protein [21]. In this approach, the experimentally obtained IR spectrum is reconstructed by non-linear fitting. For this purpose, Voigt line profiles featuring both Gaussian and Lorentzian character are mostly employed. Prior to the fitting procedure, the baseline might be subtracted. Alternatively, a baseline can be included in the fitting routine. Parameters for fitting include the number of components, band position, full width at half maximum (FWHM) and amplitude. Initial estimates for the number of fitting components and band positions can be obtained by evaluation of second derivative spectra. Generally, the number of components for this multiparameter fit (three parameters per component) should be kept low to prevent overfitting. Values for FWHM may vary between 15 and 35 cm^-1^. The goodness of fit can be assessed by evaluation of the residues, which should be below 5% of the amide I band maximum. After completing the fitting procedure, the employed software provides the area corresponding to each fitted band. Finally, the relative percentage of the individual secondary structure elements is calculated through dividing the fitted separate band areas by the sum of fitted band areas.

Fig. 4 displays an example for curve fitting of an inclusion body and folded protein spectrum. For both spectra, the same parameters for band position and FWHM were used. Table 2 shows the relative amounts of protein secondary structure determined by curve fitting. As one might expect, during protein refolding the relative amount of β-sheet decreased while α-helical content increased.

**Fig. 4.**
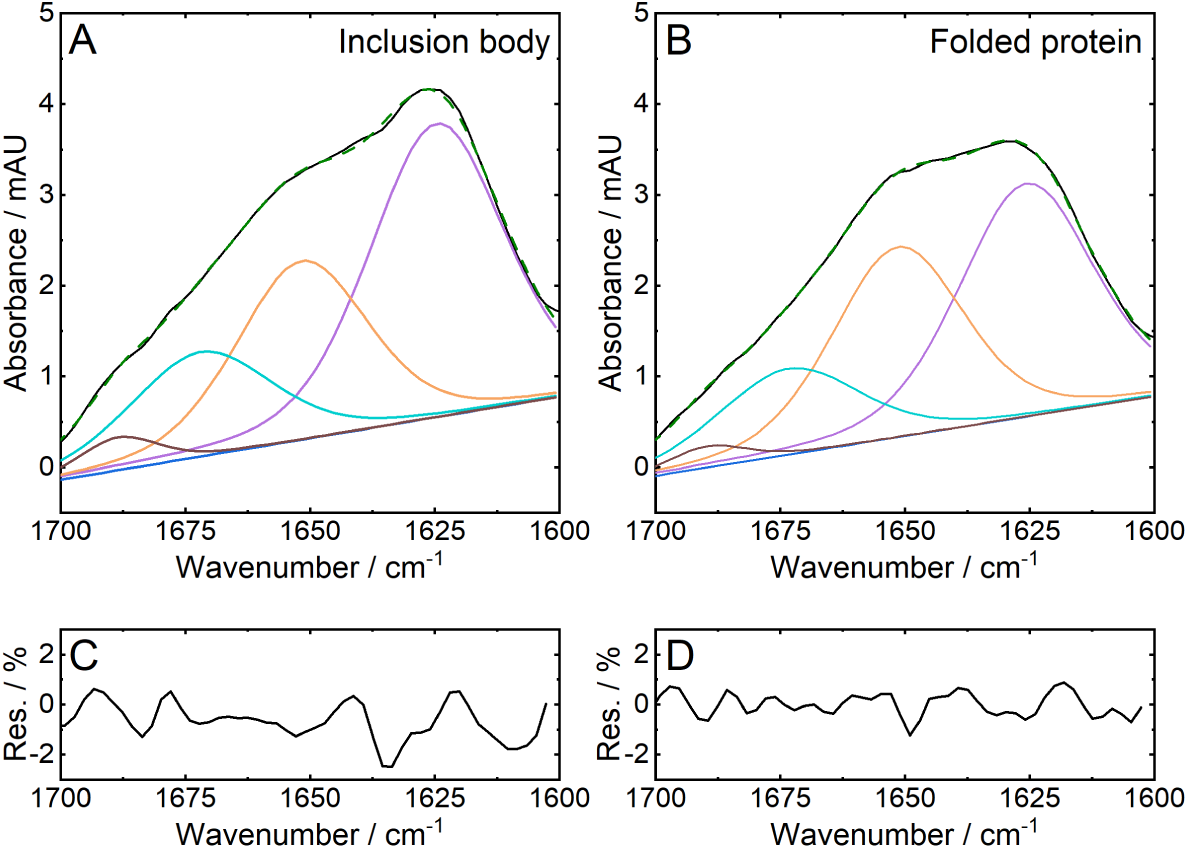
Curve fitting results of an **(A)** inclusion body and **(B)** folded protein sample with their respective residuals **(C, D)**. Residuals are calculated as percentage of the maximum band absorbance.

**Table 2.** Relative amount of secondary structure elements determined by curve fitting.

| Band<br>Maximum<br>/ $\text{cm}^{-1}$ | Assignment | Inclusion<br>Body | Folded<br>protein |
| --- | --- | --- | --- |
|  |  | Percentage / % |  |
| 1624.6 | β-sheet | 49.2 | 44.7 |
| 1651.6 | α-helix | 29.1 | 35.5 |
| 1672.0 | Turns | 18.5 | 17.4 |
| 1688.4 | β-sheet | 3.3 | 2.4 |

## 4. Notes

1. A reference absorption spectrum of water vapor (see Section 3.2.1) can be obtained by first recording a background spectrum of the empty sampling interface (i.e., ATR crystal, transmission cell) after thoroughly purging the spectrometer and the sample compartment with dry air or nitrogen. Subsequently, the flow rate of the purging system is reduced or, if possible, the sample compartment opened and the sample spectrum is recorded.
2. Noisy IR spectra can be improved by smoothing. Most software packages include a Savitzky-Golay smoothing filter. Suitable parameters are “2^nd^ polynomial order” with a window size between nine and twenty-five data points. When using large window sizes, care must be taken not to oversmooth actual spectral features.
3. IR spectra acquired in the ATR mode are not directly comparable with transmission spectra as their shape, band position and relative intensity can differ. The most striking difference is the relative band intensity distortion resulting from the wavenumber dependence of the penetration depth of the IR beam. In a transmission measurement, the path length is defined by the spacer in the flow cell and is therefore constant across the spectrum. In an ATR experiment, the depth to which the sample is penetrated by the IR beam depends on the wavelength and therefore changes throughout the spectrum. Therefore, with increasing wavenumber, the intensity of the bands decreases. Furthermore, due to anomalous dispersion, an increase in the absorption at one side of the absorption band will lead to a shift of the band maximum to lower wavenumbers [22]. In case direct comparison between IR and transmission spectra is desired, the ATR spectra should be treated with an ATR correction algorithm, which is part of most instrument software packages.
4. Air bubbles can occur more frequently in transmission flow cells with short path length. They impair the quality of IR spectra because of fringing in the individual single channel spectra, dispersion of the IR beam or incorrect absorbance values due to lack of sample. One must pay attention to remove all bubbles from the syringe before the sample is injected. In case the bubbles persist, the cell can be flushed with ethanol. It will clean the window surface and its lower surface tension increases the wettability. Subsequently, the cell should be flushed thoroughly with H_2_O or buffer.
5. The exact path length of a transmission flow cell can be calculated from a transmission spectrum of the unfilled sample cell, with the empty IR beam path as the reference. The resulting sinusoidal spectral shape, which is called interference fringe, originates from constructive and destructive interference of the IR beam from the parallel surfaces of the cell windows [23]. Using this effect, the path length of the liquid cell can be calculated by using the equation

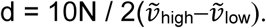
6. Here, d is the path length of the cell in millimeters, N is number of fringes in the wavenumber region spanned between 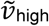 and 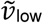. To count the number of fringes, select either minima or maxima as 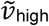 and 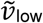 and count the minima/maxima of the spectrum in-between, as illustrated in Fig. 5. For accurate determination of the cell path length, N should be higher than five. Alternatively, most instrument software packages include automated functions for calculating the cell path length based on this approach.

**Fig. 5.**
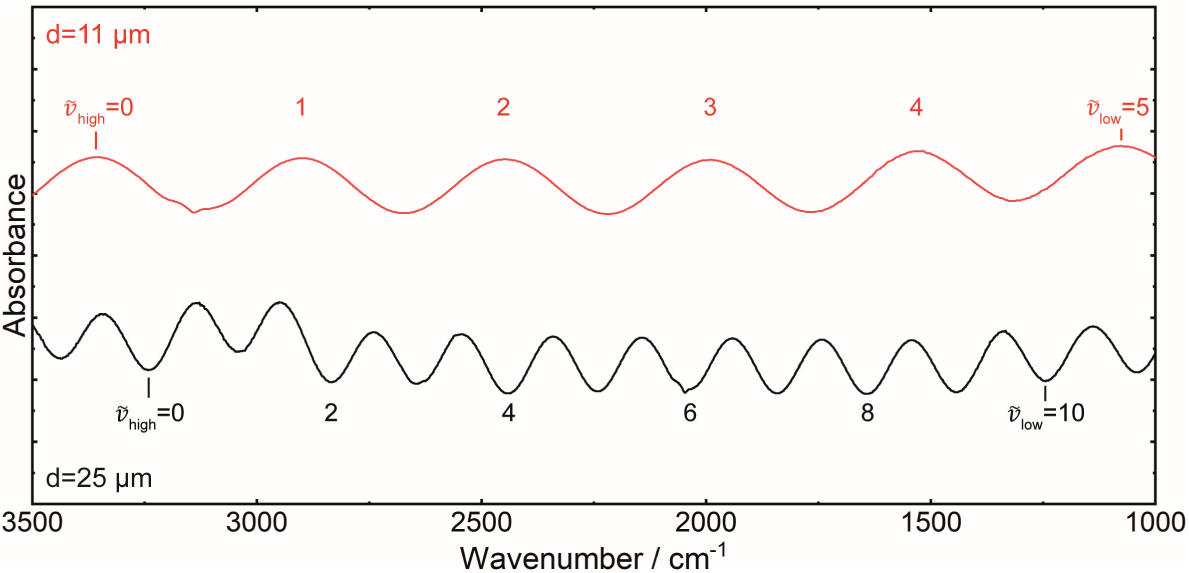
Exemplification of determination of the transmission flow cell path length.

## Acknowledgement

The authors acknowledge financial support through the Austrian Science Fund FWF (project no. P32644-N) and the COMET Centre CHASE, funded within the COMET - Competence Centers for Excellent Technologies programme by the BMK, the BMDW and the Federal Provinces of Upper Austria and Vienna. The COMET programme is managed by the Austrian Research Promotion Agency (FFG).

## References

1. Fabian H, Mäntele W. Infrared Spectroscopy of Proteins. Handbook of Vibrational Spectroscopy. Hoboken, NJ, USA: John Wiley & Sons, Ltd; 2006.

2. Yang H, Yang S, Kong J, Dong A, Yu S. Obtaining information about protein secondary structures in aqueous solution using Fourier transform IR spectroscopy. Nat Protoc. 2015;10(3):382–96. doi: 10.1038/nprot.2015.024.

3. Pelton JT, McLean LR. Spectroscopic Methods for Analysis of Protein Secondary Structure. Anal Biochem. 2000;277(2):167–76. doi: 10.1006/abio.1999.4320.

4. Barth A. Infrared spectroscopy of proteins. Biochimica et Biophysica Acta, Bioenergetics. 2007;1767:1073–101. doi: 10.1016/j.bbabio.2007.06.004.

5. de Jongh HHJ, Goormaghtigh E, Ruysschaert J-M. The Different Molar Absorptivities of the Secondary Structure Types in the Amide I Region: An Attenuated Total Reflection Infrared Study on Globular Proteins. Anal Biochem. 1996;242(1):95–103. doi: http://dx.doi.org/10.1006/abio.1996.0434.

6. Ami D, Natalello A, Taylor G, Tonon G, Maria Doglia S. Structural analysis of protein inclusion bodies by Fourier transform infrared microspectroscopy. Biochim Biophys Acta, Proteins Proteomics. 2006;1764(4):793–9. doi: http://dx.doi.org/10.1016/j.bbapap.2005.12.005.

7. Shivu B, Seshadri S, Li J, Oberg KA, Uversky VN, Fink AL. Distinct beta-sheet structure in protein aggregates determined by ATR-FTIR spectroscopy. Biochemistry. 2013;52(31):5176–83. doi: 10.1021/bi400625v.

8. Slouka C, Kopp J, Hutwimmer S, Strahammer M, Strohmer D, Eitenberger E, et al. Custom made inclusion bodies: impact of classical process parameters and physiological parameters on inclusion body quality attributes. Microbial Cell Factories. 2018;17:148. doi: 10.1186/s12934-018-0997-5.

9. Wurm DJ, Quehenberger J, Mildner J, Eggenreich B, Slouka C, Schwaighofer A, et al. Teaching an old pET new tricks: tuning of inclusion body formation and properties by a mixed feed system in E. coli. Applied Microbiology and Biotechnology. 2018;102(2):667–76. doi: 10.1007/s00253-017-8641-6.

10. Singh A, Upadhyay V, Singh A, Panda AK. Structure-Function Relationship of Inclusion Bodies of a Multimeric Protein. Frontiers in Microbiology. 2020;11. doi: 10.3389/fmicb.2020.00876.

11. Oberg K, Chrunyk BA, Wetzel R, Fink AL. Native-like Secondary Structure in Interleukin-1.beta. Inclusion Bodies by Attenuated Total Reflectance FTIR. Biochemistry. 1994;33(9):2628–34. doi: 10.1021/bi00175a035.

12. Gross-Selbeck S, Margreiter G, Obinger C, Bayer K. Fast Quantification of Recombinant Protein Inclusion Bodies within Intact Cells by FT-IR Spectroscopy. Biotechnology Progress. 2007;23(3):762–6. doi: 10.1021/bp070022q.

13. Ami D, Natalello A, Gatti-Lafranconi P, Lotti M, Doglia SM. Kinetics of inclusion body formation studied in intact cells by FT-IR spectroscopy. Febs Lett. 2005;579(16):3433–6. doi: http://dx.doi.org/10.1016/j.febslet.2005.04.085.

14. Walther C, Mayer S, Jungbauer A, Durauer A. Getting ready for PAT: Scale up and inline monitoring of protein refolding of Npro fusion proteins. Process Biochem. 2014;49(7):1113–21. doi: 10.1016/j.procbio.2014.03.022.

15. Schwaighofer A, Ablasser S, Lux L, Kopp J, Herwig C, Spadiut O, et al. Production of Active Recombinant Hyaluronidase Inclusion Bodies from Apis mellifera in E. coli Bl21(DE3) and characterization by FT-IR Spectroscopy. International Journal of Molecular Sciences. 2020;21(11):3881. doi: 10.3390/ijms21113881.

16. Mirabella FM. Internal reflection spectroscopy : theory and applications. New York; Basel: M. Dekker; 1993.

17. Akhgar CK, Ramer G, Zbik M, Trajnerowicz A, Pawluczyk J, Schwaighofer A, et al. The Next Generation of IR Spectroscopy: EC-QCL-Based Mid-IR Transmission Spectroscopy of Proteins with Balanced Detection. Anal Chem. 2020;92(14):9901–7. doi: 10.1021/acs.analchem.0c01406.

18. Schwaighofer A, Akhgar CK, Lendl B. Broadband laser-based mid-IR spectroscopy for analysis of proteins and monitoring of enzyme activity. Spectrochimica Acta Part A: Molecular and Biomolecular Spectroscopy. 2021:119563. doi: https://doi.org/10.1016/j.saa.2021.119563.

19. Goormaghtigh E, Gasper R, Bénard A, Goldsztein A, Raussens V. Protein secondary structure content in solution, films and tissues: Redundancy and complementarity of the information content in circular dichroism, transmission and ATR FTIR spectra. Biochim Biophys Acta, Proteins Proteomics. 2009;1794(9):1332–43. doi: http://dx.doi.org/10.1016/j.bbapap.2009.06.007.

20. Grdadolnik J. Infrared difference spectroscopy: Part I. Interpretation of the difference spectrum. Vib Spectrosc. 2003;31(2):279–88. doi: https://doi.org/10.1016/S0924-2031(03)00018-3.

21. Baldassarre M, Li C, Eremina N, Goormaghtigh E, Barth A. Simultaneous Fitting of Absorption Spectra and Their Second Derivatives for an Improved Analysis of Protein Infrared Spectra. Molecules. 2015;20(7):12599. doi: 10.3390/molecules200712599.

22. Ramer G, Lendl B. Attenuated Total Reflection Fourier Transform Infrared Spectroscopy. Handbook of Vibrational Spectroscopy. Hoboken, NJ, USA: John Wiley & Sons, Ltd; 2013.

23. Griffiths PR, de Haseth JA. Fourier Transform Infrared Spectrometry. Hoboken, NJ, USA: John Wiley & Sons, Inc.; 2006.

